# Crowding-induced phase separation of nuclear transport receptors in FG nucleoporin assemblies

**DOI:** 10.1101/2021.07.24.453634

**Authors:** Luke K. Davis, Ian J. Ford, Bart W. Hoogenboom

## Abstract

The rapid (< 1 ms) transport of biological material to and from the cell nucleus is regulated by the nuclear pore complex (NPC). At the core of the NPC is a permeability barrier consisting of intrinsically disordered Phe-Gly (FG) nucleoporins (FG Nups). Various types of nuclear transport receptors (NTRs) facilitate transport by partitioning in the FG Nup assembly, overcoming the barrier by their affinity to the FG Nups, and comprise a significant fraction of proteins in the NPC barrier. In previous work, we revealed that the experimental binding of the NTRs NTF2 and – the larger – Imp*β* to different planar assemblies of FG Nups follows a universal physical law defined by negative cooperativity, which was further validated by a minimal physical model that treated the FG Nups as flexible homopolymers and the NTRs as uniformly cohesive spheres ***Zahn et al. (2016***). Here, we build upon our original study by first parametrizing our model to experimental data, and next to predict the effects of crowding by different types of NTRs. We show how varying the amounts of one type of NTR modulates how the other NTR penetrates the FG Nup assembly. Notably, at similar and physiologically relevant NTR concentrations, our model predicts demixed phases of NTF2 and Imp*β* within the FG Nup assembly. The functional implication of NTR phase separation is that NPCs may sustain separate transport pathways that are determined by inter-NTR competition.

## Introduction

Embedded in the nuclear envelope are nuclear pore complexes (NPCs), large hour-glass shaped channels (inner diameter ∼ 40 nm) that regulate biomolecular transport between the cytoplasm and nucleoplasm ***Alberts (1994); Wente (2000***). The NPC presents an exclusion barrier to inert molecules, with the degree of exclusion increasing with molecular size. This physical barrier arises from a dense (mass density 100-300 mg/ml) assembly of moderately cohesive intrinsically disordered Phenylalanine-Glycine nucleoproteins (FG Nups) ***Ghavami et al. (2014***). In addition, the barrier contains relatively high numbers (∼ 20–100) of nuclear transport receptors (NTRs), globular proteins that facilitate the translocation of cargo by transiently binding to the FG Nups ***Lowe et al. (2015); Kim et al. (2018); Hayama et al. (2018***). Despite the known roles that NTRs have in nucleocytoplasmic transport, for instance ferrying specific cargo in and/or out of the nucleoplasm, returning RanGTP to the cytoplasm, and enhancing the exclusion of inert molecules ***Jovanovic-Talisman et al. (2009); Aitchison and Rout (2012); Lowe et al. (2015); Jovanovic-Talisman and Zilman (2017); Kapinos et al. (2017***), it remains to be fully elucidated how different NTRs organize themselves within the permeability barrier itself and how this organization affects transport ***Stanley et al. (2017); Jovanovic-Talisman and Zilman (2017); Hoogenboom et al. (2021***).

On the one hand, increasing the amount of different NTRs improves the performance of the nucleocytoplasmic machinery, on the other hand NTRs occupy volume which could lead to jamming in the channel. One possible explanation for how the NPC solves this problem is by modulation of the FG Nup transport barrier by NTRs or by the “separate transport pathway” hypothesis, according to which NTRs and/or cargoes may take different trajectories through the barrier, which could be determined through the differential binding of specific FG Nup – NTR pairings or spatial segregation of material in the channel ***Shah and Forbes (1998); Yang and Musser (2006); Naim et al. (2007); Fiserova et al. (2010); Yamada et al. (2010); Ma et al. (2012); Kapinos et al. (2014); Lowe et al. (2015); Lim et al. (2015); Ma et al. (2016); Kapinos et al. (2017***). An alternative mechanism, involving the switching between import and export transport states, has also been proposed ***Kapon et al. (2008***). Despite these proposals, and other candidates, there is, as yet, no definitive consensus on how the NPC maintains high-throughput transport in the presence of large numbers of NTRs ***Hoogenboom et al. (2021***).

It is difficult to test different hypotheses regarding how NTR crowding affects the NPC barrier in an *in-vivo* setting, due to the complexity of probing the intact NPC where a multitude of diverse proteins are present in a dense nanoscale channel. To circumvent this complexity, various studies have reverted to much simpler *in-vitro* FG Nup and NTR assemblies that resemble the physical environment of the NPC. Particularly well-controlled model systems are FG Nup polymer film assemblies, where copies of an FG Nup are anchored to a planar surface and NTRs (typically of one type) are introduced in the bulk volume above the surface ***Eisele et al. (2010); Schoch et al. (2012); Eisele et al. (2012); Kapinos et al. (2014); Zahn et al. (2016); Vovk et al. (2016***). For the behaviour of planar films containing one type of FG Nup and one type of NTR, the main findings have been: 1) that NTRs (such as NTF2 and – separately – Importin-*β*) bind to FG Nups in a rather generic way, suggesting possible universal physical principles – based on negative cooperative binding – governing their behaviour ***Vovk et al. (2016); Zahn et al. (2016***); 2) NTRs readily penetrate the FG Nup films, with only limited effects on the collective morphology of the FG Nups (little swelling or compaction) ***Eisele et al. (2010); Kapinos et al. (2014); Wagner et al. (2015); Vovk et al. (2016); Zahn et al. (2016***); 3) that such systems can replicate the basic selective mechanism in the NPC, *i*.*e*., inert proteins tend not to penetrate the collective FG Nup phases but NTRs do, consistent with *in-vivo* NPC functionality and with experiments on bulk solutions of FG Nups and NTRs (***Schmidt and Görlich, 2015***; ***Schmidt and Görlich, 2016***); 4) the number of transport proteins in the FG Nup films can vary by orders of magnitude as a function of NTR numbers in solution above the film, in a highly non-Langmuir manner, where complex many-body interactions preclude the use of simple one-to-one binding models ***Eisele et al. (2010); Schoch et al. (2012); Kapinos et al. (2014); Wagner et al. (2015); Vovk et al. (2016); Zahn et al. (2016***). With the caveat that only a subset of NTRs have been probed, investigations of planar assemblies of FG Nups and NTRs highlight the fine-tuned balance of FG Nup attachment density, FG Nup-NTR affinities, and NTR concentrations, where minor changes in this balance can lead to qualitatively different binding scenarios ***Vovk et al. (2016); Zahn et al. (2016); Stanley et al. (2017***).

Building on previous findings restricted to one-type of NTR, one can then ask: how does the binding of a specific NTR to a planar assembly of FG Nups depend on the presence of other NTRs? Here, we aim to provide answers to this question, using physical modelling to probe how the binding of one type of NTR could be affected by other types, in a way that can be tested by currently available experimental setups for planar assemblies of FG Nups. To explore the effects of mixed NTR crowding, we model a ternary mixture containing two different NTRs and one type of FG Nup in a polymer film assembly. When modelling FG Nups and NTRs, one can take various coarse-grained approaches, for instance one can take an all-atom approach ***Miao and Schulten (2009); Gamini et al. (2014); Raveh et al. (2016***), or account only for the amino acids (***Ghavami et al., 2012, 2014, 2018***), or work only with the generic patterning of FG/hydrophobic/hydrophilic/charged “patches” ***Tagliazucchi and Szleifer (2015); Davis et al. (2021***), or completely neglect sequence details altogether in an “homopolymer” approach ***Moussavi-Baygi et al. (2011); Osmanović et al. (2012***, 2013b); ***Vovk et al. (2016); Zahn et al. (2016); Timney et al. (2016); Davis et al. (2020***). Each approach has its strengths and weaknesses. For instance higher resolution modelling can account for greater molecular complexity, but with the difficulty in probing large time and length scales in contrast to homopolymer modelling where, at the expense of resolution, large time and length scales are accessible in conjunction with more robust parameterization and simplicity of execution. In this work, we build on our previous minimal modelling framework based on treating FG Nups as sticky and flexible homopolymers and NTRs as uniformly cohesive spheres ***Osmanović et al. (2013b); Zahn et al. (2016); Davis et al. (2020***).

## Computational Model

As in previous works ***Osmanović et al. (2013b); Zahn et al. (2016); Davis et al. (2020***), we use classical density functional theory (DFT), an equilibrium mean field theory, to model the FG Nup-NTR planar film assembly. In this work we focus solely on the effects of mixing different types of NTRs in an FG Nup film assembly, the simplest scenario being that of two types of NTRs and one type of FG Nup. Specifically, the model consists of a ternary mixture, *i*.*e*., a *ν*-component system with *ν* = 3, containing two types of free particles (NTRs denoted by *i* = 1, 2) and one type of polymer (FG Nups denoted by *i* = 3). In this system, the numbers of the two different types of free particles (components *i* = 1, 2) are given as *N*^(*i*)^, diameters are *d*^(*i*)^, and chemical potentials are *μ*^(*i*)^. In addition to the free particles, there are *N*^(i=3)^ = 260 flexible homopolymers each consisting of *M* = 300 beads, where each bead has a diameter of *d*^(3)^ =0.76 nm (corresponding to two amino acids per bead). This choice of *M* and *d*^(3)^ produces the approximate contour and persistence length of an Nsp1 FG domain ***Lim et al. (2006); Zahn et al. (2016); Hayama et al. (2019); Davis et al. (2020***). The polymers are attached uniformly to a flat surface of area *A* = 88.62 × 88.62 nm^2^, resulting in an attachment/grafting density of ≈ 3.3 polymers/100 nm^2^, which is in line with the densities in the native NPC and in *in-vitro* experiments ***Zahn et al. (2016); Davis et al. (2020***). It is assumed that the system is translationally symmetric along the directions parallel to the grafting surface, resulting in a 1D DFT theory where the densities only vary as a function of the height *z* above the surface ***Davis et al. (2020***).

Furthermore, the grand potential free energy functional Ω that provides a complete thermodynamic description of the entire system can be written as a sum of terms

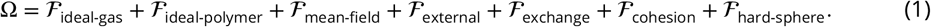

The term 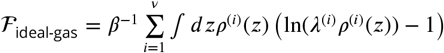 is the free energy of a set of *ν* ideal gases, where *β* = 1/*k*_B_*T* (*k*_B_ is Boltzmann’s constant) and *λ*^(*i*)^ is the appropriate thermal de Broglie wavelength for component *i*. The term ℱ_ideal-polymer_ = *N*^(3)^*β*^−1^ ln(*Z*_*c*_[*w*(*z*)]) describes the ideal polymer free energy (in the presence of a mean field *w*(*z*)) where the canonical partition function is written as

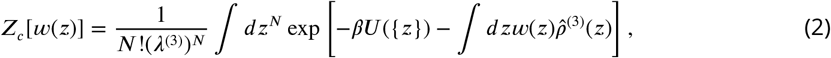

where *N* = *M* × *N*^(3)^, {*z*} is the set of all positions, *U*({*z*}) is the total potential energy, and 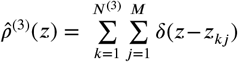 is a microscopic polymer bead density (where *z*_*kj*_ is the height of the *j*th bead belonging to polymer *k*). ℱ_mean-field_ is the additional term from introducing a polymer mean field, ℱ_external_ accounts for the external potential imposing the hardness of the anchoring surface, ℱ_exchange_ accounts for the exchange of NTRs with an external reservoir, ℱ_cohesion_ is the free energy contribution from inter- and intra-molecular attractive (“cohesive”) interactions, and ℱ_hard-sphere_ accounts for the inter- and intra-molecular excluded volume interactions, including imposing polymer chain connectivity. Thus, the dimensionless grand potential can be written more explicitly as

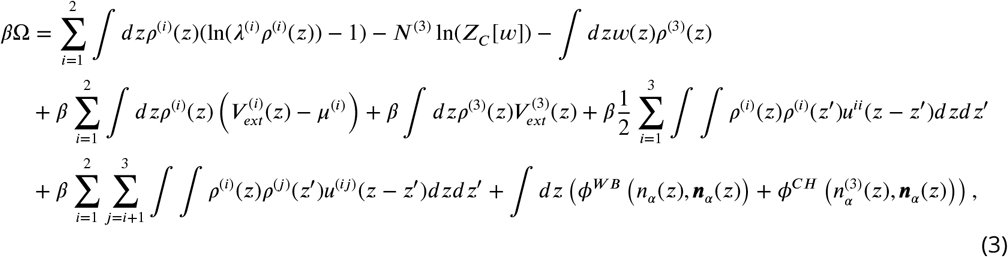

where *ρ*^(*i*)^(*z*) is the number density profile of component *i*, 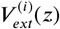 is the external potential representing the surface, *μ*^(*i*)^ is the chemical potential for component *i, u*^*ij*^ (*z*) is a one-dimensional pair potential defined between components *i* and *j, ϕ*^*WB*^ and *ϕ*^*CH*^ are the White bear (hard-sphere) ***Roth et al. (2002***) and chain connectivity functionals ***Yu and Wu (2002***) given by the equations

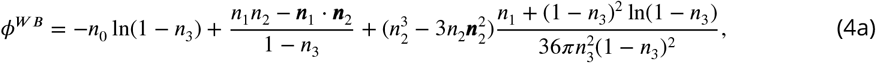

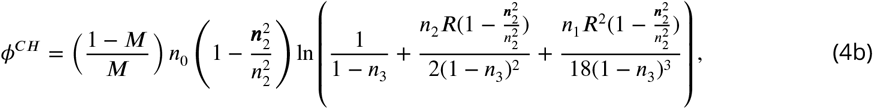

where *n*_*α*_(*z*; {*ρ*^(*i*)^}) and *n*_*α*_(*z*; {*ρ*^(*i*)^}) are, respectively, the one-dimensional scalar and vectorial weighted densities and *R* is the radius of a polymer bead (see ***Roth (2010***) and ***Davis et al. (2020***)).

The hardness of the flat surface is imposed via a Weeks-Chandler-Anderson potential

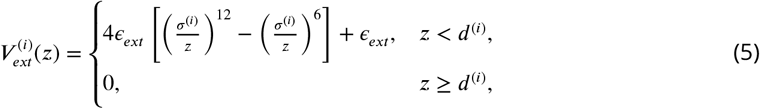

where *ϵ*_*ext*_ = 20 *k*_B_*T* is the maximum energy barrier of the wall (chosen sufficiently high so that the number density of all components is zero at and below the surface) and *σ*^(*i*)^ = 2^−1/6^*d*^(*i*)^. Consistent with our previous work ***Davis et al. (2021***), the intramolecular and intermolecular cohesive interactions are based upon the Morse potential (in three dimensions)

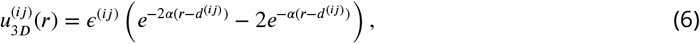

where *r* is the distance between two particles of type *i* and type *j, ϵ*^(*ij*)^ is the cohesion strength, *α* = 6.0 nm^−1^ is an inverse decay length of the pair potential, and 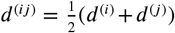. The potential above, valid in three spatial dimensions, is then integrated over the plane, henceforth only depending on *z*, and shifted such that – the now one-dimensional pair potential – *u*^(*ij*)^(*z* ≥ 2*d*^(*ij*)^) = 0 *k*_B_*T* so as to keep the cohesive interactions short ranged.

To find the set of density distributions – for the particles and polymer – and the polymer mean field in the equilibrium state, the following equations must be solved self-consistently

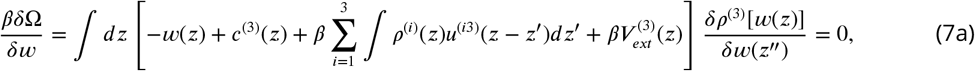

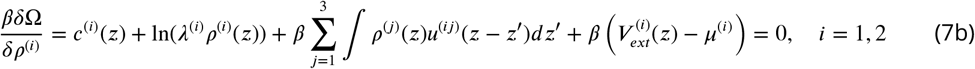

where the notation 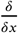 represents a functional derivative with respect to *x* and *c*^(*i*)^ is the one-body direct correlation function given by

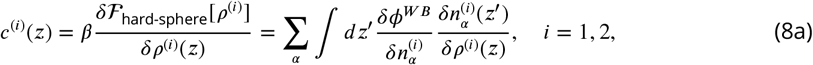

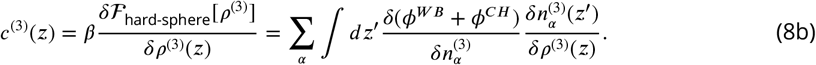

For the free particles one can decompose the chemical potential into two terms

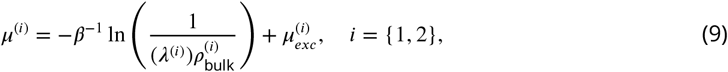

where 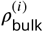 is the bulk density of the free particles of component *i* and 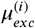 is the excess chemical potential due to the inter- and intra-molecular interactions. One can solve equations (7) self-consistently by invoking a fictitious time variable *t*, where the solutions are found through an iterative procedure. This is expressed by the following

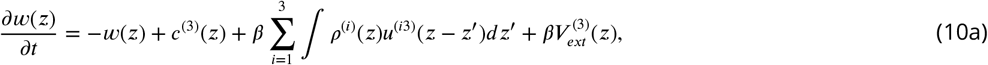

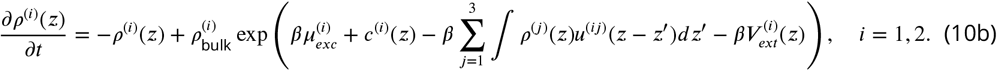

Finally, discretizing space *z* into *L* slices of thickness Δ*z* and discretizing fictitious time then yields the resulting discrete update rules which are solved numerically

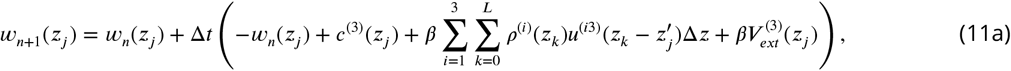

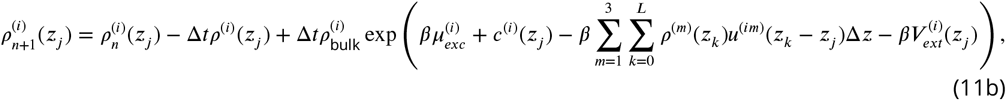

where *z*_*k*_ is the – midpoint – height above the surface of the spatial slice *k, n* labels discrete time, and in the last equation *i* = 1, 2. The simulation parameters used here were *L* = 1024, Δ*z* = 0.117 nm (with *z*_0_ = 0.0585), and Δ*t* = 0.002.

## Results

Minimal physical modelling facilitates the understanding of many aspects of NPC functionality in terms of general principles, but it requires quantitatively accurate parameter settings to make meaningful predictions ***Osmanovic et al. (2013a); Jovanovic-Talisman and Zilman (2017); Hoogenboom et al. (2021***). In this work the minimal modelling framework we employ is that of coarse-grained classical density functional theory (DFT), which has been previously used to model aspects of the NPC permeability barrier ***Osmanović et al. (2012***, 2013b); ***Zahn et al. (2016); Davis et al. (2020***). To ensure that the setting of the parameters in our DFT model – outlined above – is commensurate with the behaviour of FG Nups and NTRs as probed in experiments, we make use of experimental data on FG Nup-NTR polymer film assemblies where the macroscopic binding between one type of FG Nup and one type of NTR was quantitatively probed (see Figure 1) ***Zahn et al. (2016***). The experiments focussed on a polymer film consisting of FG Nup (Nsp1) domains, attached to a flat surface at physiologically relevant densities (≈ 3.3 polymers/100 nm^2^), interacting with one of the two following NTRs: NTF2, and Importin-*β* (Imp*β*).

**Figure 1.**
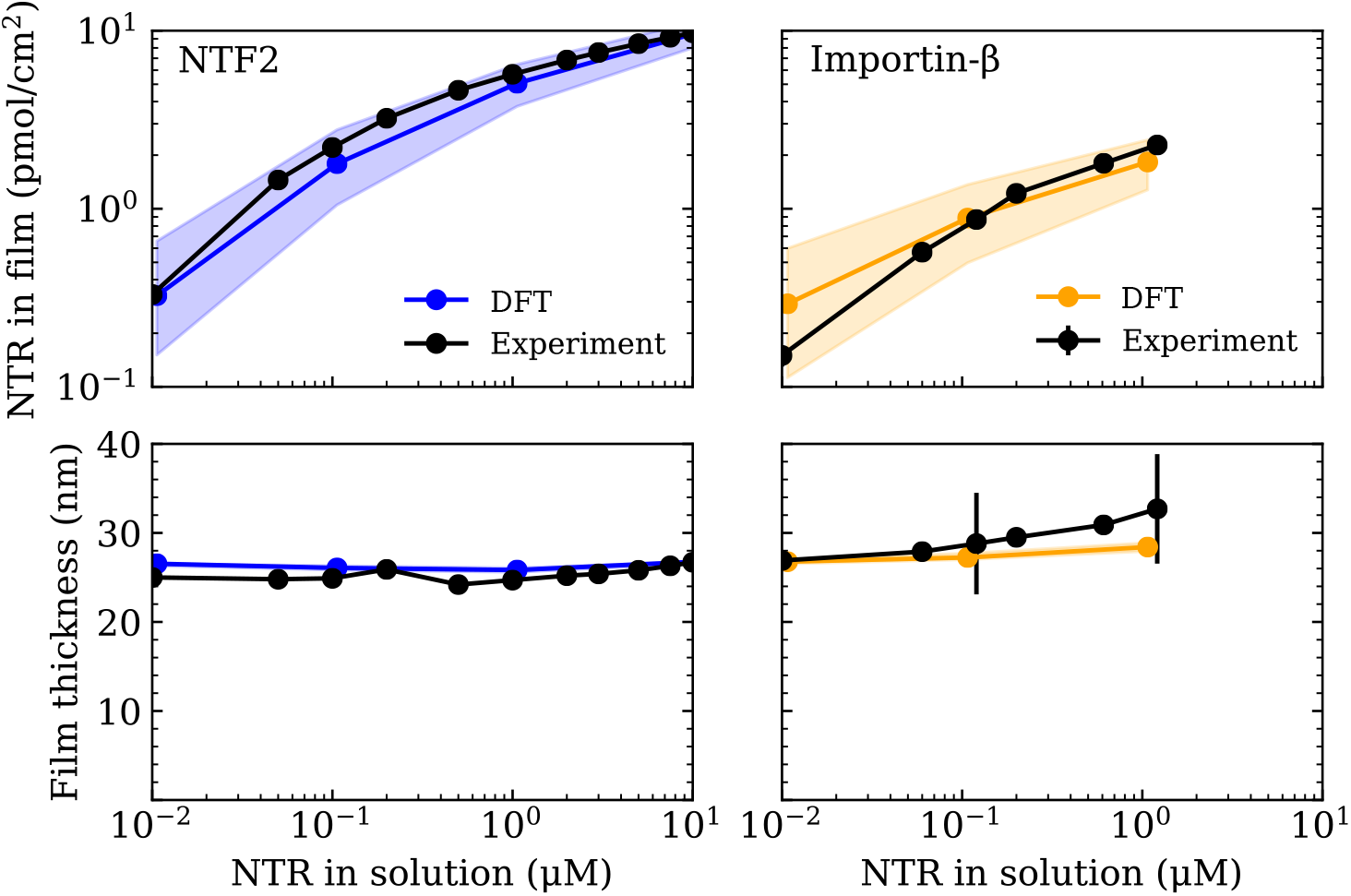
Setting the polymer-particle cohesion strengths {*ϵ*_FG-NTF2_, *ϵ*_FG-Imp*β*_} through comparison of DFT results with experimental binding isotherms for the cases of NTF2 (left) and Importin-*β* (right) binding to an Nsp1 film ***Zahn et al. (2016***) (top). Concomitant film thicknesses as found in DFT and experiment (bottom). The experimental Nsp1 surface attachment densities were 4.9 pmol/cm^2^ and 5.1 pmol/cm^2^ for NTF2 and Importin-*β* respectively. The parametrized cohesion strengths *ϵ*_FG-NTF2_ = 2.4 ± 0.1 *k*_B_*T* and *ϵ*_FG-Imp*β*_ = 2.3 ± 0.1 *k*_B_*T* correspond to the modelled NTF2 and Importin-*β* particles respectively. Filled bands (in all four panels) denote a tolerance of ±0.1 *k*_B_*T* in the polymer-particle cohesion strengths. The thicknesses of the filled bands for the bottom two panels is similar to the thickness of the line connecting DFT data points (blue and orange). **Figure 1–Figure supplement 1**. Parameterizing the polymer cohesion strength *ϵ*_FG-FG_. **Figure 1–Figure supplement 2**. Inert particles of growing size do not penetrate the polymer film.

For consistency with the available experimental data we focus on the FG Nup Nsp1, which we treat as a homogeneous, flexible, and cohesive polymer consisting of *M* = 300 beads of diameter 0.76 nm (2 amino acids per bead), resulting in the approximately correct persistence length for FG Nups ***Lim et al. (2006); Zahn et al. (2016); Hayama et al. (2019); Davis et al. (2020***). The inter- and intra-molecular cohesive properties of FG Nups arise from a combination of hydrophobic motifs, *e*.*g*., FG, FxFG, and GLFG, and charged residues along the sequence which, in our model, are subsumed into one essential cohesion parameter *ϵ*_FG-FG_. In addition to the FG Nups, we also include the presence of the NTRs NTF2 and Imp*β*, which we model as uniformly cohesive spheres of diameters *d*_NTF2_ = 4 nm and *d*_Imp*β*_ = 6 nm respectively ***Zahn et al. (2016***). The cohesive properties of the NTRs strictly refer to the attraction between the NTRs and FG Nups, arising at least in part from the hydrophobic grooves and charged regions on the former and the hydrophobic motifs and charged regions on the latter ***Kumeta et al. (2012); Kim et al. (2013); Hayama et al. (2018); Frey et al. (2018***). Following previous work ***Zahn et al. (2016); Vovk et al. (2016***), we do not consider any cohesion between NTRs themselves. As with the FG Nup inter- and intra-molecular cohesive interactions, we subsume all contributions to the respective cohesive interactions FG Nup – NTF2 and FG Nup – Imp*β* through two more cohesion parameters *ϵ*_FG-NTF2_ and *ϵ*_FG-Imp*β*_.

We begin the parametrization of our model with the setting of *ϵ*_FG-FG_ so as to accurately reproduce the experimental thickness of Nsp1 planar film assemblies, at similar anchoring densities, as was done previously ***Zahn et al. (2016); Fisher et al. (2018); Davis et al. (2020***) (see Figure 1-Figure supplement 1). With the here chosen interaction potential, the resulting cohesion strength is *ϵ*_FG-FG_ = 0.275 ± 0.025 *k*_B_*T* (with experiments conducted at ≈ 23 °C), which yields a film thickness *τ* = 26 ± 2 nm, in our model defined as the height above the surface below which 95% of the total polymer density is included. We note that the value of *ϵ*_FG-FG_ found here is different to that of our previous work ***Zahn et al. (2016***), which is due to the different choices of interaction potential and geometry of the film assembly, but both models are parametrized using the same experimental data and produce the same film thicknesses. To further validate this value of *ϵ*_FG-FG_, we checked whether the polymer film would exclude inert molecules, a basic property that has been observed for Nsp1 assemblies (***Schmidt and Görlich, 2015***; ***Schmidt and Görlich, 2016***). The inert molecules are modelled as non-cohesive spheres of diameter *d*^(*i*)^, with *i* labelling the particle type, and their inclusion/exclusion in the film is quantified through the potential of mean force (PMF) *W* ^(*i*)^(*z*) given as

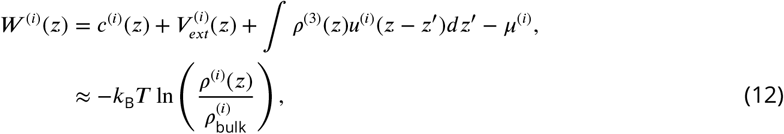

where *i* denotes a particle type, *c*^(*i*)^(*z*) is the one-body direct correlation function (see equation 8), 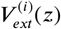 is the external potential (see equation 5), *ρ*^(3)^(*z*) is the polymer number density, *u*^(*i*)^(*z*) is the one-dimensional (integrated over the *x* − *y* plane) polymer-particle cohesive pair potential (see equation 6), and 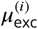 is the excess chemical potential (here set to 0 *k*_B_*T*) ***Roth et al. (2000); Roth and Kinoshita (2006***). For the inert molecules, the polymer-particle attraction (third term in equation (12)) is nullified. As expected, non-cohesive particles with increasing diameters (1.0, 2.0, 4.0, and 6.0 nm) experience a greater potential barrier upon attempted entry into the polymer film (see Figure 1-Figure supplement 2), confirming that our simple model of an Nsp1 film replicates one of the key characteristics of the permeability barrier as seen in the NPC: the degree of exclusion of inert molecules increasing with molecular size ***Mohr et al. (2009); Popken et al. (2015); Ghavami et al. (2016***). We note that the presence of a maximum, close to the anchoring surface, in the relative density for inert particle diameters *d* = 1 and 2 nm is due to the decrease in the polymer density closer to the surface (consistent with a small potential well close to the surface see Figure 1-Figure supplement 2b); the appearance of the maxima in the density depends upon the anchoring density of the FG Nups (not shown here). With the energy of thermal fluctuations *k*_B_*T* as a reference point, PMFs of the order of *k*_B_*T* are compatible with passive entry and exit whilst PMFs at least one order of magnitude greater than *k*_B_*T* are indicative of a significant barrier. Hence we find that inert particles with diameters ⪆ 4 nm experience a large free energy barrier to penetrate the FG Nup assembly. This is quantitatively similar to previous experimental estimates of the “soft” size selectivity *d* = 4.5 − 5.4 nm of the NPC ***Keminer and Peters (1999); Paine et al. (1975); Mohr et al. (2009***), and consistent with another simulation study which explicitly accounted for the amino acid sequence of the FG Nups ***Ghavami et al. (2016***).

Having shown that the now parametrized polymer model for Nsp1 films replicates the experimentally observed film thickness and the size selectivity of the NPC, we shift our focus to setting the parameters for the NTRs NTF2 and Imp*β*. The cohesion strengths *ϵ*_FG-NTF2_ and *ϵ*_FG-Imp*β*_ are set by comparing absorption isotherms as calculated in DFT to those measured in experiment ***Zahn et al. (2016***) (see Figure 1). Using DFT, we compute the total density of NTRs in the film Γ^(*i*)^, *i* = {1, 2}, as the total NTR population within the film thickness *τ* divided by the area *A* (converted to units of pmol/cm^2^)

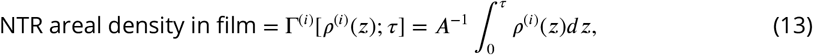

where *ρ*^(*i*)^(*z*) (*i* = {1, 2}) is the number density of the NTRs. With only one free fitting parameter for each NTR (for the NTR-FG Nup interaction strength), the DFT binding isotherms are found in excellent agreement with experiment over 3 orders of magnitude in bulk NTR concentration (Figure 1 (top)), as was previously accomplished by a similar DFT model (where polymers were attached to the base of a cylinder) in ***Zahn et al. (2016***). The resulting parametrized cohesion strengths – for the here chosen interaction potential – are *ϵ*_FG-NTF2_ = 2.4 ± 0.1 *k*_B_*T* and *ϵ*_FG-Imp*β*_ = 2.3 ± 0.1 *k*_B_*T* for NTF2 and Imp*β* respectively. One might notice that *ϵ*_FG-NTF2_ ≈ *ϵ*_FG-Imp*β*_ for the two (model) NTRs, despite the Imp*β* particle having an 1.5-fold larger excluded-volume diameter as compared with the NTF2 particle. However, given the differences in diameters, and therefore a difference in the respective ranges of intermolecular interactions (see equation 6), we caution against directly comparing the cohesive properties of the two NTRs based on the cohesion strengths *ϵ*_FG-NTF2_ and *ϵ*_FG-Imp*β*_ alone. Of note, the concomitant film thicknesses from DFT – as a function of NTR concentration – are also in good agreement with experiment (Figure 1 (bottom)).

At this point, all the essential interaction parameters *ϵ*_FG-FG_, *ϵ*_FG-NTF2_, and *ϵ*_FG-Imp*β*_ have been set by quantitative comparisons between DFT and experiment. Next, we investigate the effects of crowding of one type of NTR on the system. We quantify molecular crowding through two observables: (i) the packing fraction *ρ*^(*i*)^(*z*)*d*^(*i*)^, where *ρ*^(*i*)^(*z*) is the one-dimensional number density of a particular NTR (labeled by *i*), and (ii) the potential of mean force (PMF) *W* ^(*i*)^(*z*) of a particular NTR, in the presence of other NTRs and the FG Nups (see Figure 2 and equation 12). For high crowding, one expects the packing fraction of a particular NTR to increase in magnitude and for the PMF to become more positive (with respect to the same system but with fewer NTRs), which is interpreted as a greater potential barrier (or, somewhat equivalently, a shallower potential well). We observe that both NTF2 and Imp*β* display higher levels of packing and higher-amplitude density oscillations within the Nsp1 film upon increasing their respective bulk concentrations (0.01, 0.1, 1.0, and 10.0 μM) (Figure 2 (top)). The density oscillations arise from layering/ordering effects mainly caused by packing against a hard planar wall, where particles prefer to pack closer to a flat surface; the layering of hard-spheres next to a planar wall is a well known phenomenon ***Patra (1999); Roth et al. (2000); Deb et al. (2011***). As is expected, in both systems, the maximum observed packing fraction (⪆ 0.15) was located close to the surface (at 10 μM). For the here chosen NTR-particle sizes, it is expected that the packing fraction and PMF will be largely dictated by the interactions with the polymers and the crowding of other NTRs, with less significant effects arising from the particular implementation of the surface hardness. We note that the density oscillations for the Imp*β* particle show greater amplitudes as compared with the NTF2 particle (for the same concentration above the film), which is expected since the Imp*β* is larger in size and thus experiences more pronounced layering effects ***Padmanabhan et al. (2010***).

**Figure 2.**
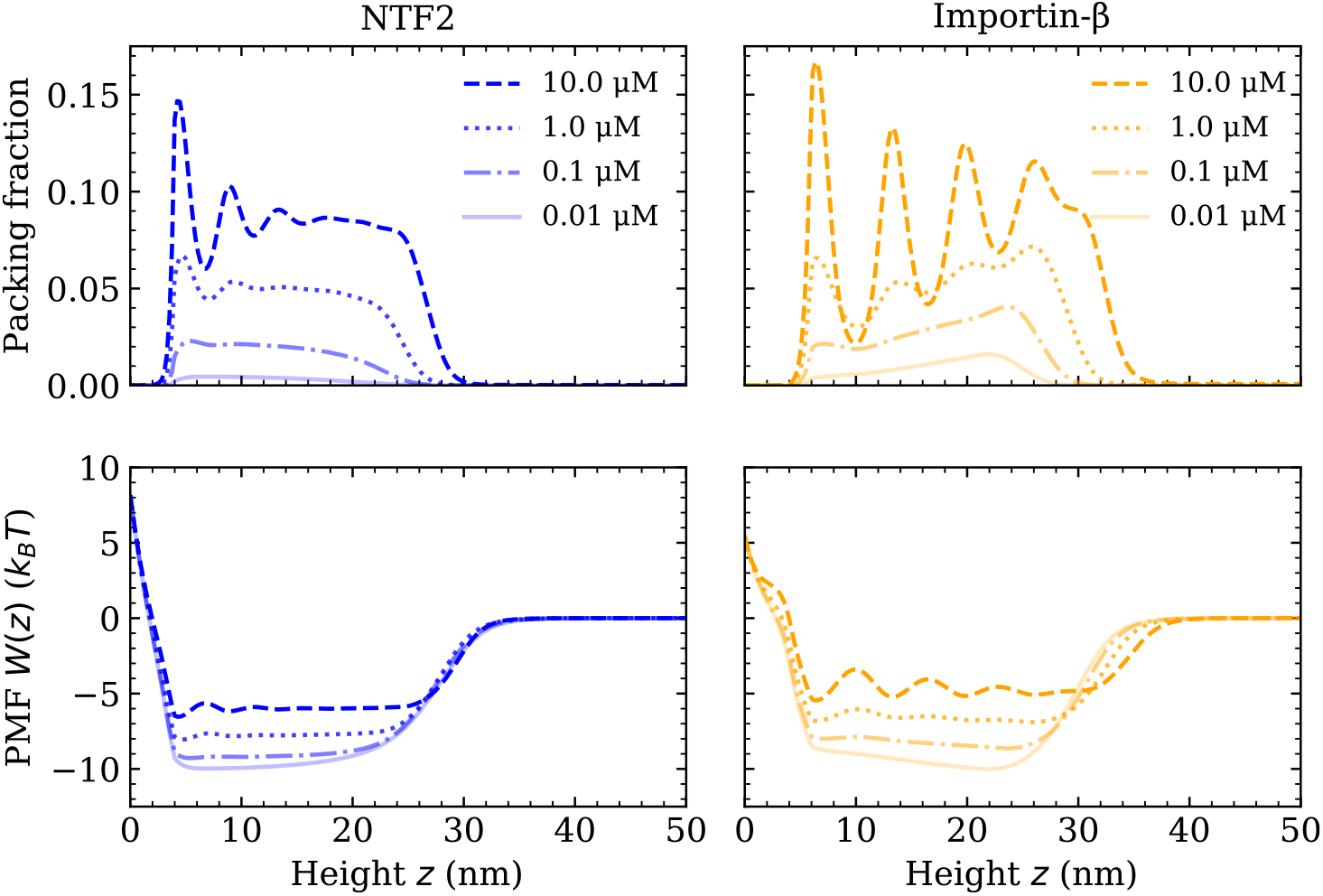
Increasing NTR bulk concentration increases packing and filling up of the potential well within the Nsp1 film, for systems containing one type of NTR only. Equilibrium DFT packing fractions *ρ*(*z*)*d*, where *ρ*(*z*) is the one-dimensional number density and *d* is the particle diameter, as a function of the height *z* above the flat surface for NTF2 (left) and Importin-*β* (right), at various concentrations (top). Accompanying potentials of mean force *W* (*z*) (bottom), for the same systems as on the top row.

For both NTF2 and Imp*β*, the PMFs decrease in magnitude (but remain negative within the bulk of the film) upon an increase in bulk NTR concentration (Figure 2 (bottom)). Specifically, increasing the concentration in solution from 0.01 to 10.0 μM results in an approximate two fold decrease in the absolute value of the PMF (|Δ*W* (*z*)| ≈ 4 − 5 *k*_B_*T*). The implication of this finding is that, at higher levels of packing in the film, it is relatively easier for bound NTRs to unbind from the polymer film, or, equivalently, less favourable for additional NTRs to enter the polymer film from the solution above it. This effect may primarily be attributed to the increased filling of space, *i*.*e*. molecular crowding, of the NTRs between the Nsp1 polymers. The results of Figure 2 are particularly relevant to the NPC “transport paradox”, where fast transport (∼ 1000 transport events per second) occurs in conjunction with strong – selective – binding. Whilst there are various explanations of the transport paradox ***Hoogenboom et al. (2021***), these results show how NTR crowding may facilitate the exit of NTRs from the NPC, noting that a decrease of |Δ*W* (*z*)| ≈ 4 − 5 *k*_B_*T* in PMF well depth would imply a ∼100× faster rate for unbinding if we assume Arrhenius-like kinetics (Figure 2 (bottom)).

As a next step, we explore how the competition between NTRs may affect the binding, penetration, and distribution of NTRs in FG Nup assemblies. Specifically, we model the mixed crowding effects in a system containing the NTRs NTF2 and Imp*β* in an Nsp1 polymer film (see Figures 3 and 4 and their respective Figure supplements). As in the case with one type of NTR, we probe the packing fractions, potentials of mean force (PMFs) *W* ^(*i*)^(*z*), binding isotherms, and polymer film thickness but this time keeping the amount of one NTR fixed at a physiologically relevant concentration (1 μM) ***Zahn et al. (2016***) whilst varying the concentration of the other NTR (Figure 3a). Upon increasing the bulk concentration of NTF2 from 0.01 μM to 10.0 μM while keeping the bulk concentration (in solution) of Imp*β* constant at 1 μM, the amount of bound Imp*β* dramatically drops and the remaining bound Imp*β* is redistributed towards the surface of the Nsp1 polymer film (Figure 3a (top) and Figure 3-Figure supplement 1). Additionally, the density oscillations of Imp*β* within the film, which are evident at 0.01 μM of NTF2, smooth out upon increasing the amount of NTF2 to 0.1 μM. This shows that the presence of NTF2 directly modulates the distribution of Imp*β* within the film. Interestingly, upon increasing NTF2 from 0.1 μM while keeping the bulk concentration of Imp*β* constant, we observe NTR phase separation: an NTF2-rich phase within the FG Nup film and an Imp*β*-rich phase at the film surface.

**Figure 3.**
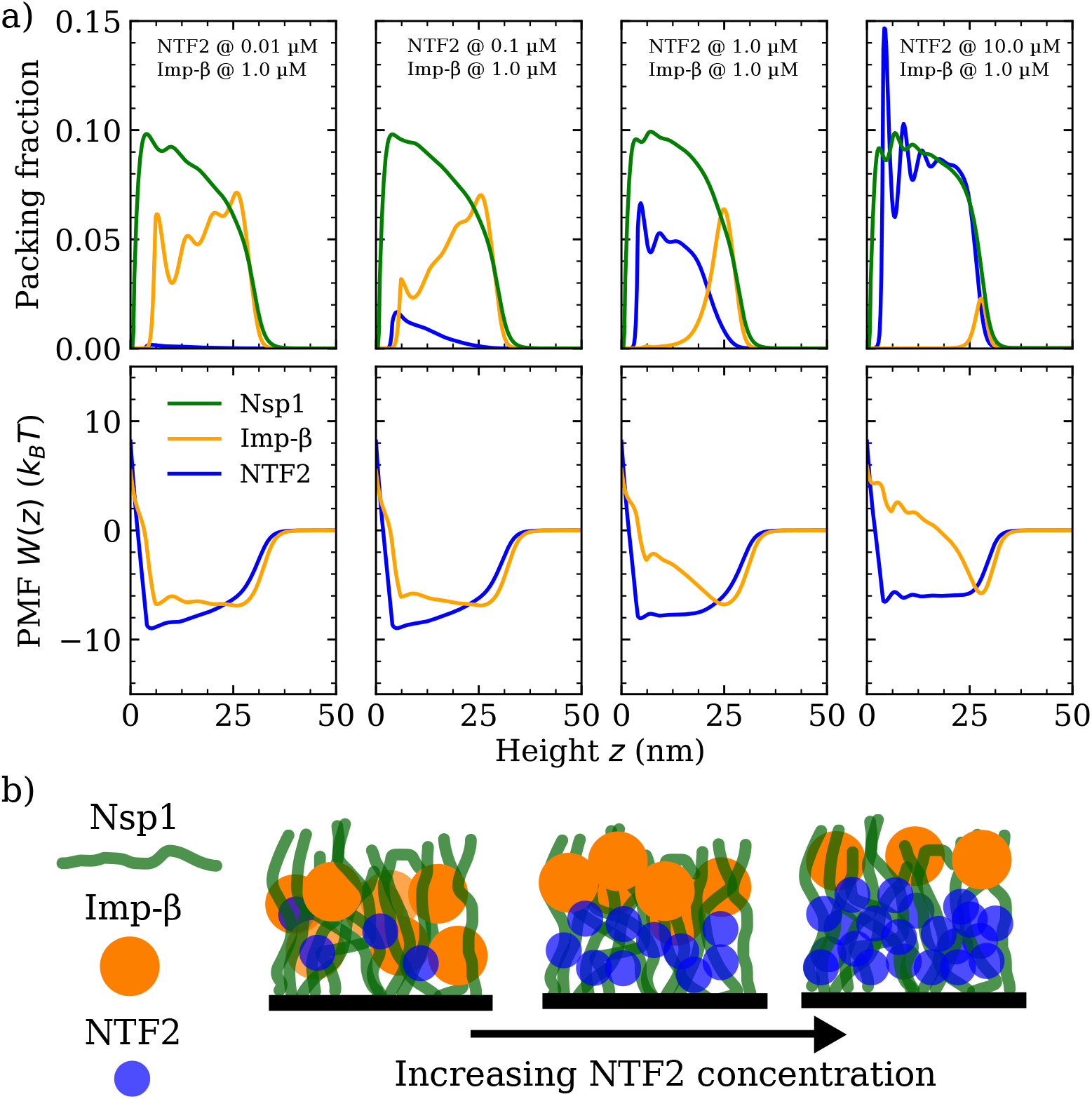
Phase separation in a ternary FG Nup-NTR polymer film assembly. **a)** DFT Packing fractions and accompanying potentials of mean force (PMFs) for Nsp1 polymer films in the presence of NTF2 and Importin-*β* (Imp*β*). The concentration (in solution) of NTF2 is increased from 0.01-10.0 μM (left to right panels), whilst the concentration of Imp*β* is fixed at 1.0 μM. The cohesion strengths used here are {*ϵ*_FG-FG_ = 0.275, *ϵ*_FG-NTF2_ = 2.4, *ϵ*_FG-Imp*β*_ = 2.3} *k*_B_*T* for the Nsp1-Nsp1, Nsp1-NTF2, and Nsp1 - Imp*β* interactions respectively. **b)** Cartoon visualisation of the data from (a) depicting the increasing concentration of NTF2 pushing Imp*β* to the top of the Nsp1 layer, also resulting in significant expulsion of Imp*β* from the film into the bulk. **Figure 3–Figure supplement 1**. NTR binding isotherms and Nsp1 film thicknesses as a function of NTF2 concentration in solution.

**Figure 4.**
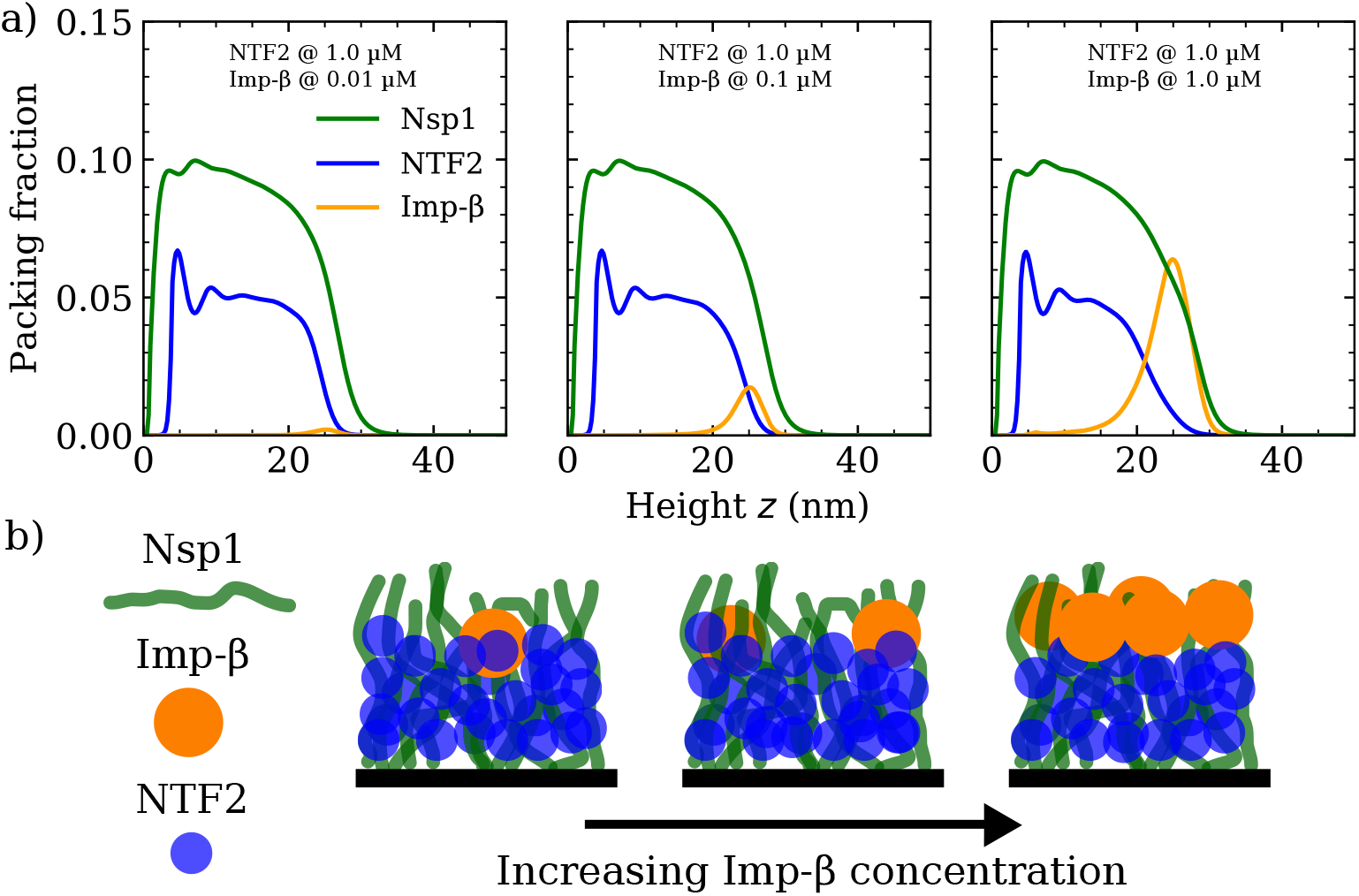
Increasing Importin-*β* (Imp*β*) concentration negligibly affects NTF2 in the Nsp1 film. **a)** DFT packing fractions against the height above the flat surface *z* for Nsp1, NTF2, and Imp*β*. The concentration of Imp*β* is increased from 0.01-10 μM (left to right panels) whilst the NTF2 concentration remains fixed at 1.0 μM. The last panel (furthest to the right) is the same as the second last panel in Figure 3a. **b)** Cartoon illustration visualising the data in (a) depicting the undetectable change in the packing/morphology of the NTF2 in the presence of increasing Imp*β* molecules. **Figure 4–Figure supplement 1**. NTR binding isotherms and Nsp1 film thicknesses as a function of Importin-*β* concentration in solution.

When considering binary systems of hard-spheres with different diameters subject to packing between planar walls, ignoring any attractive interactions between them, one typically observes the larger particles packing closer to the wall, as compared with the smaller particles ***Padmanabhan et al. (2010***). This effect, as measured per unit area, is due to the overall system entropy loss being less when the larger particles pack closer to the surface, rather than the smaller ones. Here we observe the opposite effect, with the (smaller) NTF2 particles remaining closer to the grafting surface, which is qualitatively consistent with a theoretical study investigating a binary mixture of attractive particles, where the larger particles were excluded for a distance from a planar surface of up to twice the particle diameter ***Padmanabhan et al. (2010***). Here, however, we observe the depletion of the larger NTR (Imp*β*) over much larger distances (in *z*) for high bulk concentrations of NTF2, apparently dictated by the polymer film thickness. The intuitive explanation is that the smaller NTF2 competes more readily for binding sites (that are spread uniformly along the polymer in our model) deep within the film, closer to the grafting surface, without paying a substantial entropic penalty for rearranging the polymers. In contrast, closer to the film surface, the larger Imp*β* binds more readily since its overall stronger binding propensity (note *ϵ*_FG-NTF2_ ≈ *ϵ*_FG-Imp*β*_, spread over a larger particle surface for Imp*β*) where the polymers are more diffuse. Indeed, the distribution of NTF2 in the film largely follows the polymer density as a function of distance from the grafting surface, indicating that with its smaller size, NTF2 benefits more from the enhanced concentration of polymer beads (and therewith of binding sites) without having to pay a substantial entropic cost (as for Imp*β*) for penetrating the polymer film.

Throughout the changes in incorporation of NTF2, the density of the polymers did not show noticeable changes. The modulation of Imp*β* via changes in NTF2 numbers is also articulated in terms of the PMF *W* (*z*), where the Imp*β* PMF is an approximate square well at 0.01 μM of NTF2, but for higher NTF2 concentrations gradually transforms into a pronounced and sharper well located at *z* ≈ 25.0 nm, *i*.*e*., at the surface of the film, with the formation of a barrier to enter the rest of the film (Figure 3a (bottom)).

Finally, we verified if similar effects resulted when increasing the concentration of Imp*β* for a given, constant, NTF2 concentration set at 1.0 μM (see Figure 4a and Figure 4-Figure supplement 1). We observe no significant change to the Nsp1 or NTF2 packing factions (including the PMF and binding isotherm) upon increasing the concentration of Imp*β* in solution from 0.01-1.0 μM (see also Figure 4b). We have not explored high bulk concentrations (> 1 μM) of Imp*β*, as these yield highly oscillatory packing fractions and therewith are computationally more challenging in our DFT model. However, we expect that further incorporation of Imp*β* would eventually change the distribution of NTF2 in the film.

## Discussion

We have made quantitative predictions regarding the effects of mixed crowding on the binding of different NTRs to FG Nup planar assemblies. Our results are based on a minimal coarse-grained model implemented in a mean-field theory approach, which treats FG Nups as sticky and flexible homopolymers and NTRs as isotropic and cohesive spheres. Firstly, the model includes three interaction parameters, corresponding to the cohesive interactions between FG Nups and NTRs: these were parametrized using experimental data for Nsp1 film assemblies and binding thereto of one type of NTR (NTF2 or Imp*β*).

Based on the thus parametrized model, we have shown that increased crowding of one type of NTR results in a shallower potential well within the film, implying that individual NTRs will have a small potential barrier to leaving the film in the presence of more NTRs. The origin of this effect is due to an interplay between the further occupation of volume within the film (entropic) and the increased competition for binding sites. This result has important implications for the NPC: when there is a large influx of material into the channel from either the cytoplasm or nucleoplasm, the exit of said material should be faster since the increased crowding effects will reduce the free energy barrier – thus increasing the likelihood – to leave the pore, with a predicted increase in unbinding rates of two orders of magnitude in the concentration range explored here. While we note that there are multiple factors involved in determining transport speed ***Hoogenboom et al. (2021***), this scenario highlights the importance of NTRs as an essential component in the NPC transport barrier ***Lim et al. (2015***) and, specifically, implies that the NPC could perform more efficiently and faster with higher numbers of NTRs present in its inner channel, as has indeed been observed in experiments with Imp*β* ***Yang and Musser (2006***).

We found that with increased incorporation of NTF2 within the FG Nup film, the amount of absorbed Imp*β* was reduced and its distribution within the film was changed. For similar and physiologically relevant concentrations of the NTRs studied here, phase separation was observed, with an NTF2-rich phase at the bottom of the film (where the polymer packing is higher) and an Imp*β*-rich phase at the top of the film (where the polymer packing is lower). It is important to note that this particular height dependent phase separation, as predicted in our minimal one dimensional model (assuming symmetry parallel to the anchoring surface), might not be the only way NTRs spatially segregate. Therefore, it would be interesting to extend the model developed here to two or three dimensions, relaxing the lateral symmetry assumption (see ***Osmanović et al. (2013b***)). Additionally, future developments of our approach could explore the implications that mixed NTR crowding has on kinetics, with careful considerations of how one coarse-grains the sequence heterogeneity of FG Nups and the patchiness of the NTRs as this is important for kinetics ***Davis et al. (2021***).

Given that there is a stable population of Imp*β* in the NPC barrier and given that changes to this population affect the selective properties of the NPC ***Lowe et al. (2015***), our results suggest that NTR crowding plays a substantial role in determining the performance of the NPC barrier. Additionally, the observation of a phase-separated state between two distinct NTRs has implications on how the NPC maintains high-throughput transport despite high NTR densities. Consistent with experimental observations on NPCs ***Lowe et al. (2015***), Imp*β* is found to occupy regions of lower FG Nup density (as shown here in planar FG Nup assemblies), where our results here demonstrate that such a distribution of Imp*β* can at least in part be attributed to competitive binding of other, smaller NTRs to regions of higher FG Nup density. Future work could explore mixed crowding in the pore geometry of the NPC where the FG Nup density decreases with a growing radial distance away from the center of the pore, giving rise to a possible – radially dependent – phase separation in the NPC. An immediate consequence of this is that transport pathways through the NPC are most likely dependent on the type of NTR, with potentially separate transport pathways mediated and modulated by different NTRs.

## Acknowledgments

We thank Dino Osmanović, Anton Zilman, and Andela Šaric for discussions. We thank Ralf Richter for providing feedback on the manuscript. L.K.D. acknowledges the biophysics research computing cluster at UCL that was used to perform the simulations and analysis. This work was funded by the UK Engineering and Physical Sciences Research Council (EP/L504889/1, L.K.D. and B.W.H.).

---

**Figure 1–Figure supplement 1.**
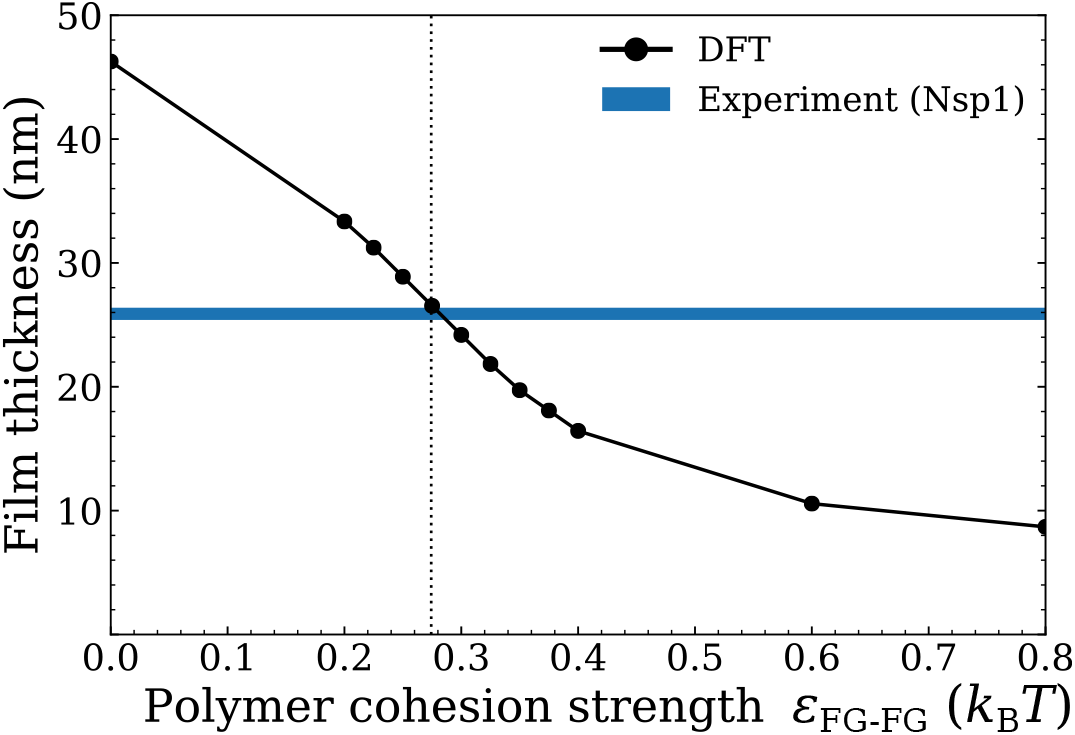
Setting the polymer-polymer cohesion parameter *ϵ*_FG-FG_ through comparison of film thicknesses as calculated from DFT, *i*.*e*., the height including 95% of polymer density, with the film thickness of an Nsp1 film assembly as derived from experiment (25.9±0.5 nm) ***Zahn et al. (2016***). The grafting density of Nsp1 to the flat surface in DFT was set so as to best match the density used in experiments (4.9 pmol/cm^2^ ≈ 3.3 polymers/100nm^2^). The vertical dotted line corresponds to the interpolated *ϵ*_FG-FG_ = 0.275 *k*_B_*T* for which the DFT best matches the experimental thickness.

**Figure 1–Figure supplement 2.**
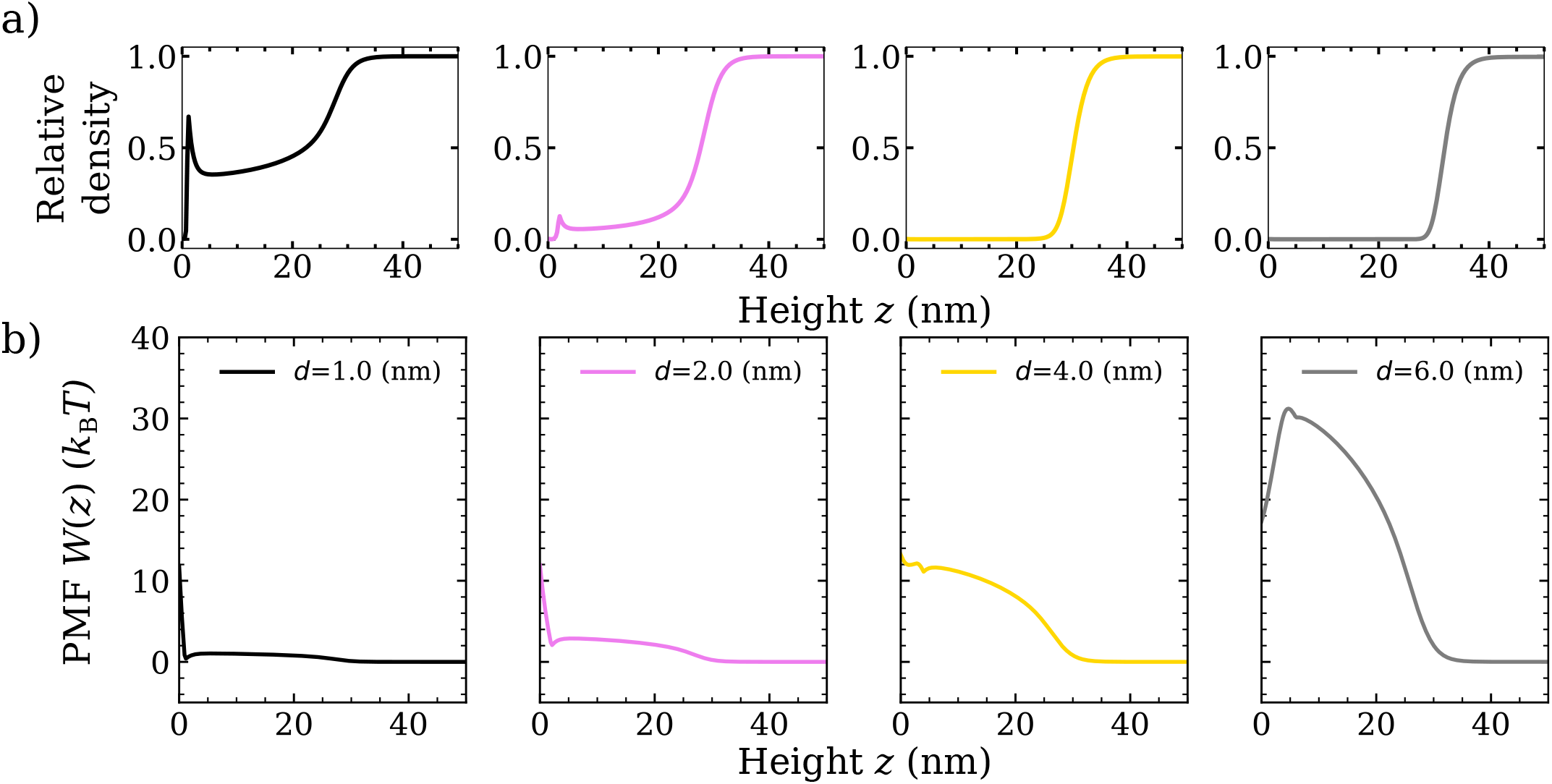
Quantification of entry (to the Nsp1 film) barriers for non-cohesive particles with varying diameters *d*. **a)** Relative density (normalized to the bulk value occurring at *z* ≥ 40 nm) of non-cohesive particles with diameters *d* = 1.0 nm (black), *d* = 2.0 nm (pink), *d* = 4.0 nm (gold), *d* = 6.0 nm (grey) as calculated in classical density functional theory (DFT). **b)** Potential of mean force (PMF) as a function of the height above the flat surface *z*. The concentration of the particles in the solution is 10.0 μM for all panels. The polymer-polymer cohesion strength is *ϵ*_FG-FG_ = 0.275 *k*_B_*T*, as set through comparison with an experimental Nsp1 film.

**Figure 3–Figure supplement 1.**
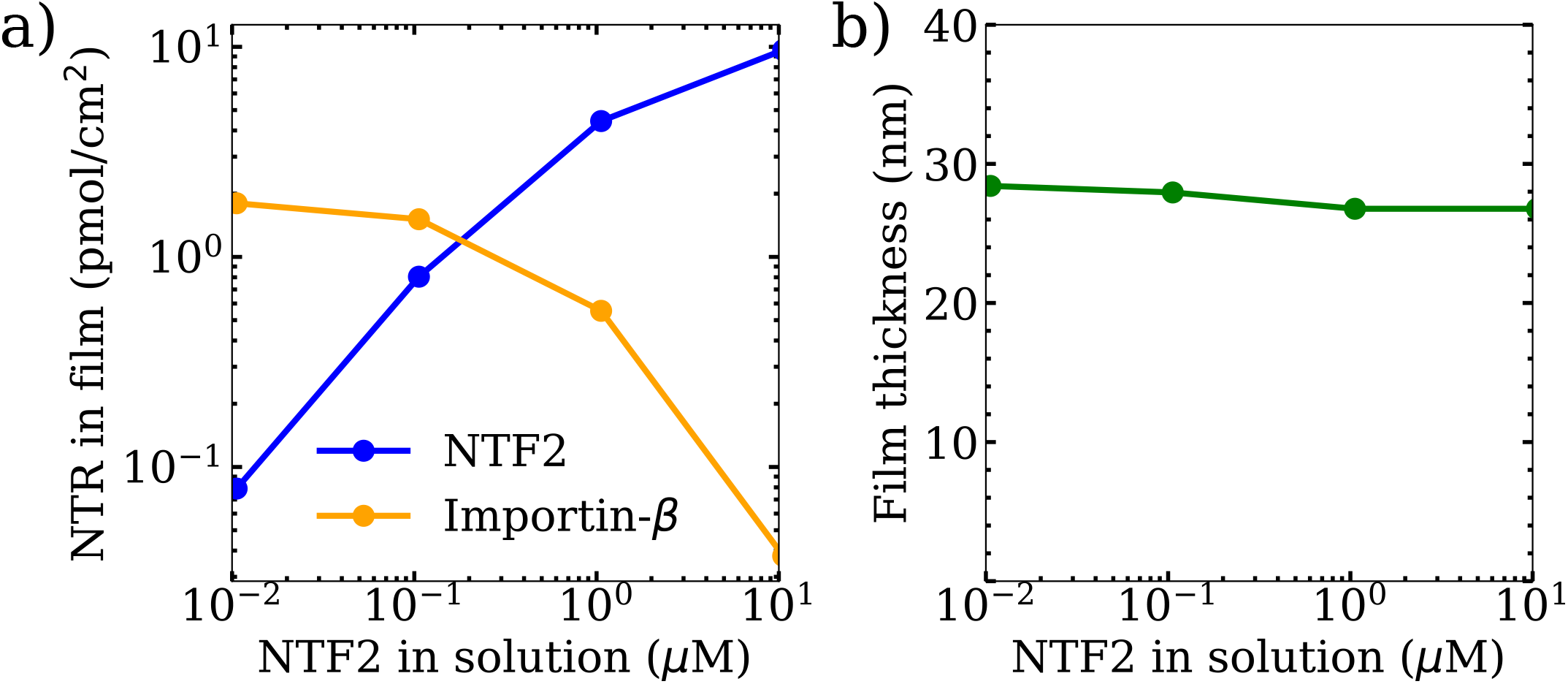
NTR binding isotherms and Nsp1 film thicknesses as a function of NTF2 concentration in solution. **a)** Binding isotherms as predicted from the classical density functional (DFT) model for NTF2 and Importin-*β*. **b)** Concomitant film thickness of the FG Nup (Nsp1) layer as found in DFT. The cohesion strengths used here are {*ϵ*_FG-FG_ = 0.275, *ϵ*_FG-NTF2_ = 2.4, *ϵ*_FG-Imp*β*_ = 2.3} *k*_B_*T* for the Nsp1-Nsp1, Nsp1-NTF2, and Nsp1 - Imp*β* interactions respectively. In each plot the concentration of Importin-*β* in solution remained – approximately – fixed at 1 μM.

**Figure 4–Figure supplement 1.**
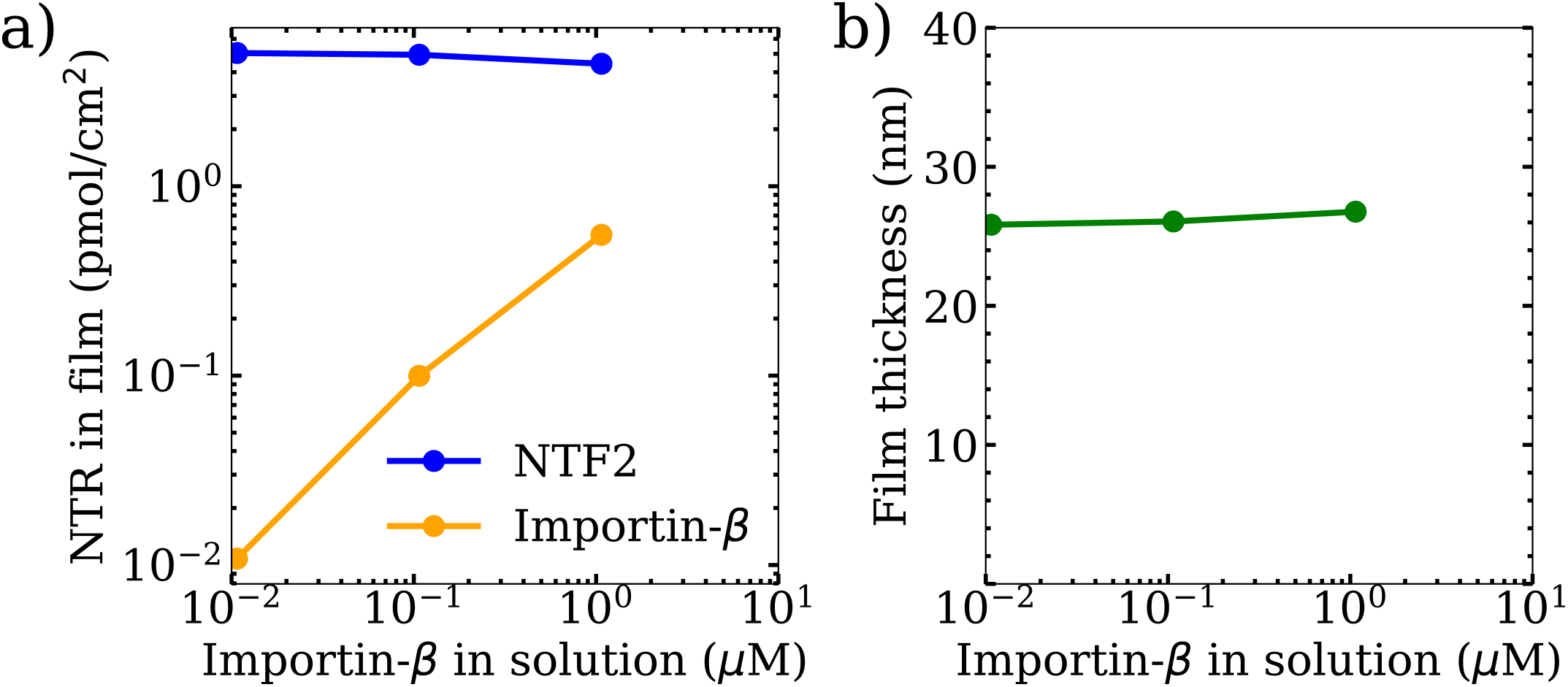
NTR binding isotherms and Nsp1 film thicknesses as a function of Importin-*β* concentration in solution. **a)** Binding isotherms as predicted from the classical density functional (DFT) model for NTF2 and Importin-*β*. **b)** Concomitant film thickness of the FG Nup (Nsp1) layer as found in DFT. The cohesion strengths used here are {*ϵ*_FG-FG_ = 0.275, *ϵ*_FG-NTF2_ = 2.4, *ϵ*_FG-Imp*β*_ = 2.3} *k*_B_*T* for the Nsp1-Nsp1, Nsp1-NTF2, and Nsp1 - Imp*β* interactions respectively. In each plot the concentration of NTF2 in solution remained – approximately – fixed at 1 μM.

## Notes

### Competing Interest Statement

The authors have declared no competing interest.

